# Emergence and diversification of a host-parasite RNA ecosystem through Darwinian evolution

**DOI:** 10.1101/728659

**Authors:** Taro Furubayashi, Kensuke Ueda, Yohsuke Bansho, Daisuke Motooka, Shota Nakamura, Norikazu Ichihashi

## Abstract

In the prebiotic evolution, molecular self-replicators are considered to develop into diverse, complex living organisms. The appearance of parasitic replicators is believed inevitable in this process. However, the role of parasitic replicators on prebiotic evolution remains elusive. Here, we demonstrated experimental coevolution of RNA self-replicators (host RNAs) and emerging parasitic replicators (parasitic RNAs) for the first time by using an RNA-protein replication system we had developed. During a long-term replication experiment, a clonal population of the host RNA turned into an evolving host-parasite ecosystem through the continuous emergence of new types of host and parasitic RNAs produced by replication errors. The diversified host and parasitic RNAs exhibited evolutionary arms-race dynamics. The parasitic RNA accumulated unique mutations that the host RNA had never acquired, thus adding a new genetic variation to the whole replicator ensemble. These results provide the first experimental evidence that the coevolutionary interplay between host-parasite molecules play a key role in generating diversity and complexity in prebiotic molecular evolution.

## Introduction

Host-parasite coevolution is at the center of the entire course of the biological evolution (Claverie, 2006; Forterre and Prangishvili, 2009; Koonin and Dolja, 2013; Koskella and Brockhurst, 2014). Parasitic replicators, such as viruses, are the most prosperous biological entities (Bergh et al., 1989; Suttle, 2007) that offer ever-changing selection pressure and genetic reservoirs in the global biosphere. The invention of the sophisticated adaptive immunity (Müller et al., 2018) prevailing all the domains of life is a hallmark to display the power of host-parasite coevolution, and accumulating evidence further highlights the potential key roles of parasites for the development of the basic biological architectures and functions (Claverie, 2006; Deininger et al., 2003; Elbarbary et al., 2016; Forterre and Prangishvili, 2013, 2009; Iranzo et al., 2014; Koonin and Dolja, 2013; Koskella and Brockhurst, 2014).

Parasitic replicators probably worked as the evolutionary drivers from the prebiological era of molecular replication (Higgs and Lehman, 2015; Joyce and Szostak, 2018; Orgel, 1992; Szathmáry and Maynard Smith, 1997; Wochner et al., 2011). Even in a simplest form of replication system, parasites inevitably appear through a functional loss of self-replicating molecules and threaten the sustainability of the replication system (Bansho et al., 2012; Koonin et al., 2017). Theoretical studies suggested that spatial structures, such as cell-like compartments, allow self-replicators (i.e. hosts) to survive by limiting the propagation of parasitic replicators (Bresch et al., 1980; Furubayashi and Ichihashi, 2018; Szathmáry and Demeter, 1987; Takeuchi and Hogeweg, 2009). Following experimental studies demonstrated that the compartmentalization strategy effectively supports the replication of the host replicators in the presence of parasitic replicators (Bansho et al., 2016, 2012; Matsumura et al., 2016; Mizuuchi and Ichihashi, 2018).

In a previous study (Ichihashi et al., 2013), for the purpose of studying how a simple molecular system develops through Darwinian evolution, we constructed an RNA replication system consisting of an artificial genomic RNA and a reconstituted translation system of *Escherichia coli* (Shimizu et al., 2001) encapsulated in water-in-oil droplets. In this system, the artificial genomic RNA (host RNA) replicates through the translation of the self-encoded replicase subunit. During replication, a deletion mutant of the host RNA (parasitic RNA), which lost the encoded replicase subunit gene, spontaneously appears and replicates by freeriding the replicase provided from the host RNA. Through serial nutrient supply and dilution, the host and parasitic RNAs in water-in-oil droplets undergo repeated error-prone replication and natural selection processes, i.e. Darwinian evolution.

In a following study (Bansho et al., 2016), to study the evolutionary process of the host and parasitic RNA replicators, we performed a serial transfer replication experiment of the RNA replication system. In this work, we reported that the host and parasitic RNAs showed oscillating population dynamics and the host RNA acquired a certain level of parasite-resistance at the final rounds of the replication experiment (43 rounds, 215 h). However, we did not observe counter-adaptative evolution of the parasitic RNA to the host RNA, and the coevolutionary process of the host and parasitic RNAs still remains unclear.

In this study, we thought that a much longer time is necessary for coevolution of the host and parasitic RNA replicators, we extended the replication experiment for additional 77 rounds (385 h). To understand their evolutionary dynamics during the replication experiment, we performed sequence analysis of the host and parasitic RNAs. We also conducted competitive replication assays using evolved host and parasitic RNA clones to confirm the coevolution between the host and parasitic RNAs. Note that we fully reanalyzed the host-parasite RNA population up to 43 rounds partially reported in the previous study (Bansho et al., 2016) and present the whole picture of the 120 rounds (600 h) of the long-term replication experiment with new data in this manuscript.

## Results

### RNA replication system

The RNA replication system used in this study consists of two classes of single-stranded RNAs (host and parasitic RNAs) and a reconstituted translation system of *E. coli* (Shimizu et al., 2001) (Fig.1A). A distinctive feature between the host and parasitic RNAs is the capability of providing an RNA replicase (Qβ replicase). The host RNA provides the catalytic β-subunit of the replicase via translation, which forms active replicase by associating with EF-Tu and EF-Ts subunits in the translation system, while the parasitic RNA lacks the intact gene. The host RNA replicates using the self-provided replicase, whereas the parasitic RNA replicates relying on the host-provided replicase. As the original host RNA, we used a clone from round 128 in our previous study (Ichihashi et al., 2013).

**Fig.1.**
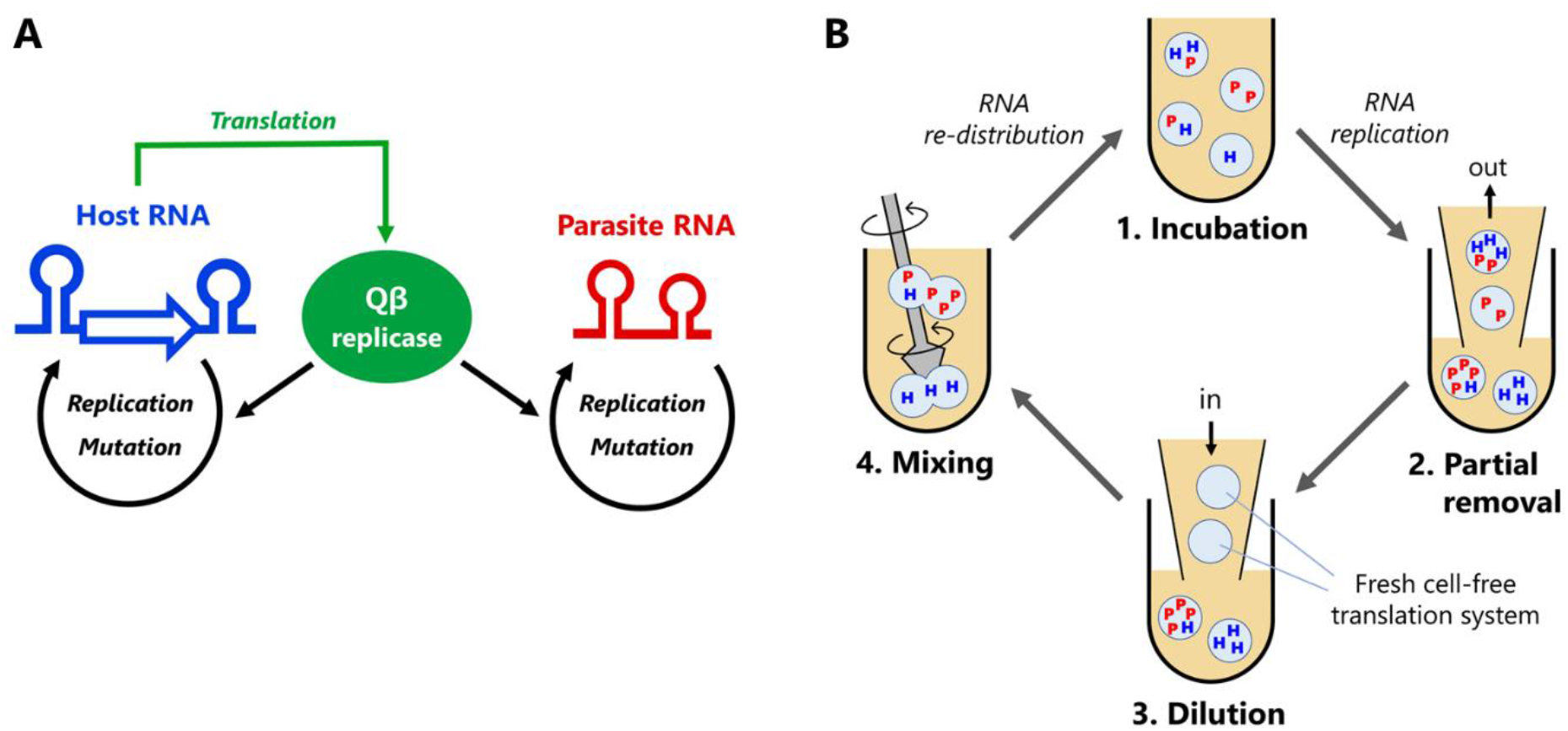
Host and parasitic RNA replication system. (A) Replication scheme of the host and parasitic RNAs. The host RNA encodes Qβ replicase subunit, while the parasitic RNA does not. Both RNAs are replicated by the translated Qβ replicase in the reconstituted translation system of *E. coli.* (B) Replication-dilution cycle for a long-term replication experiment. The host RNA is encapsulated in water-in-oil droplets with approximately 2 μm diameter. The parasitic RNA spontaneously appears. (1) The droplets are incubated at 37 °C for 5 h for translation and replication. (2) 80 % of droplets are removed and (3) diluted with new droplets containing the translation system (i.e., 5-fold dilution). (4) Diluted droplets are vigorously mixed to induce random fusion and division among the droplets. We repeated this cycle for 120 rounds. Reaction volume was 1 mL with 1% aqueous phase, corresponding to approximately 10^8^ droplets.

In this system, parasitic RNAs spontaneously emerges from the host RNA by deleting the internal replicase gene plausibly through nonhomologous recombination (Bansho et al., 2016; Chetverin et al., 1997). The parasitic RNAs reported previously have similar sizes (~200 nt). We refer to this size of parasitic RNA as “parasite-α”. The parasite-α replicates much faster than the original host RNA (~2040 nt) due to its smaller size, and thus inhibits the host replication through competition for the replicase. The replication with Qβ replicase is error-prone, about 1.0× 10^-5^ per base (García-Villada and Drake, 2012), and mutations are randomly introduced into the host and parasitic RNAs during the replication reaction.

### Long-term replication experiment

We performed a long-term replication experiment of the host and parasitic RNAs. The replication reaction was performed in water-in-oil droplets through repeating a fusion-division cycle with the supply of new droplets containing the translation system (Fig.1B). The single round of the experiment consists of four steps: 1) incubation, 2) partial removal, 3) dilution, and 4) mixing. 1) In the incubation process, the water-in-oil droplets were incubated at 37 °C for 5 h to induce the internal translation and RNA replication reactions. Note that we started with a clonal population of the host RNA (1nM, ~ 6×10^9^ molecules) without the parasite-α, but the parasite-α was detected within two rounds. 2) In the partial removal process, we randomly removed 80% of the water-in-oil droplets and 3) substituted with new water-in-oil droplets containing the cell-free translation system in the dilution process (i.e., 5-fold dilution). 4) In the mixing process, droplets were vigorously mixed with a homogenizer to induce random fusion and division among droplets to allow the mixing of RNAs and other components among droplets. We repeated this cycle for 120 rounds (600 h) in total.

### Population dynamics of the host and parasitic RNAs

We measured the concentrations of the host and parasitic RNAs after every incubation process (Fig.2A). The host RNA was measured using quantitative PCR after reverse transcription (RT-qPCR) and the parasitic RNA was measured from the band intensity after polyacrylamide gel electrophoresis (Supplementary Figure 1). In some rounds (7-12, 16-22, and 75-84), the parasitic RNAs were under the detection limit (less than ~30 nM).

**Fig.2.**
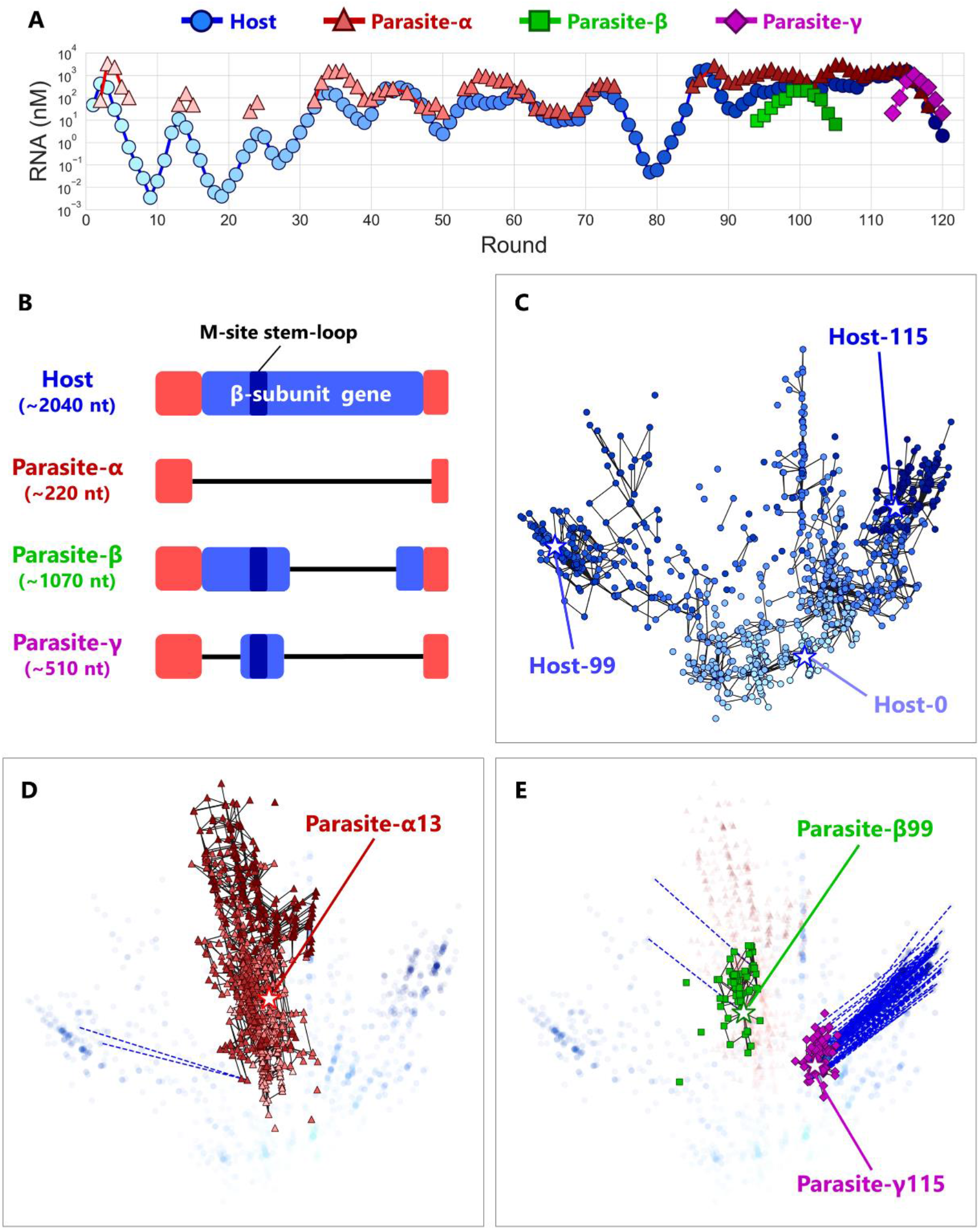
Coevolutionary dynamics of the host and parasitic RNAs. (A) Population dynamics of the host and parasitic RNAs through the long-term replication experiment. In the regions without points, parasitic RNA concentrations were under the detection limits (< 30 nM). Three different parasitic species (α, β, and γ) are classified based on their sizes. (B) Schematic representation of the sequence alignments of the host and parasitic RNA species. The terminal regions (red) of all the RNA species are derived from a small replicable RNA by the replicase, MDV-1 (Mills et al., 1973). β-subunit encoding region are shown in blue and the branched stem-loop of the M-site, one of the binding site for Qβ replicase, is also indicated. Deleted regions are shown in black lines. (C, D, E) 2D maps of the dominant RNA genotypes for the host RNA (C), parasites-α (D), -β and (E) -γ. The top-90 dominant genotypes were plotted for each round. A point represents each genotype. The color depths are consistent with those in (A). Black lines connect pairs of points one Hamming distance apart in the same RNA species. A broken blue line connects a pair of points zero Hamming distance apart (perfect match) in the different RNA species, ignoring the large deletion between host and parasitic RNAs. Stars represent the genotypes of the evolved RNAs used for the competitive replication assay in Fig.3A. The original host RNA is Host-0. Round-by-round data are shown in Supplementary Figure 5.

The population dynamics of the host and parasitic RNAs gradually changed throughout rounds (Fig.2A). In the early stage (round 1 to ~35), the host RNA and the parasite-α showed a relatively regular oscillation pattern caused by competition between the host and parasitic RNAs in compartments. In this regime, the concentrations of the parasite-α were higher than those of the host RNA where detected. In the middle stage (round ~35 to ~75), the concentrations of the host RNA increased and the oscillation pattern became irregular, suggesting that the host RNA acquired parasite resistance. In the later stage (round ~95 to ~116), the concentrations of the host and the parasite-α further increased and the oscillation pattern became further unclear. In this regime, we observed the appearance of new parasitic RNA species of different sizes and classified them as parasites-β (~1000 nt, green squares) and -γ (~500 nt, purple diamonds) according to their sizes. We termed these new RNAs “parasites” because each clone of these RNAs does not replicate alone (Supplementary Figure 2). These continuously changing population dynamics can be caused by successive adaptation processes between host and parasitic RNAs.

### Sequence analysis

To investigate evolutionary dynamics of host-parasite RNA populations at the sequence level, we recovered RNA mixtures from 17 points (rounds 13, 24, 33, 39, 43, 50, 53, 60, 65, 72, 86, 91, 94, 99, 104, 110, and 115) and subjected them to reverse transcription followed by deep sequencing with PacBio RS II for the host, parasite-β and parasite-γ and MiSeq for the parasite-α. For the sequencing using PacBio RS II, we obtained 365-4143 reads for each class of RNA in sequenced rounds. For the sequencing using MiSeq, we obtained ~5000 reads for each round (Supplementary Figure 3).

Sequence analysis revealed that four major RNA classes with different sizes existed in the long-term replication experiment, consistent with the band positions observed in the polyacrylamide gels: the host (~2040 nt), the parasite-α (~220 nt), the parasite-β (~1070 nt) and the parasite-γ (~510 nt). The sequences of all the classes of parasitic RNA shared a high degree of similarity with those of host RNAs but lacked a large part of the replicase subunit gene (Fig.2B). The parasite-α lacks the entire gene. The parasite-β lacks approximately 3’ half of the gene and the parasite-γ further lost approximately 1/4 of the remaining 5’ region of the gene. Both the parasite-β and -γ retain a part of the M-site sequence, one of the recognition sites for Qβ replicase (Meyer et al., 1981; Schuppli et al., 1998), in the middle of the gene.

We next determined dominant genotypes of all the classes of RNA (host, parasite-α, -β, and -γ). Although the RNAs replication by Qβ replicase is error-prone and introduces many random mutations to produce quasi-species for each genotype, we focused on the consensus sequences that consist of mutations commonly found in the RNA population. We first identified 74 dominant mutations that were present more than 10% of the population of each class of RNA in a sequenced round. The dominant mutations consisted of 60 base substitutions, 4 insertions, and 10 deletions in total (Supplementary Figure 4). Next, we measured the frequencies of all the genotypes composed of the combination of these 74 dominant mutations in every sequenced round for each class of RNA. All the genotypes and their frequencies are shown in Supplementary Data.

We next investigated the relationships of the detected genotypes. To visualize evolutionary trajectories, we calculated Hamming distances between all combinations of the top-90 genotypes of all the classes of RNA species in the sequenced rounds and then plotted them in a single two-dimensional (2D) map by using Principle Coordinate Analysis (Fig. 2C-E). Note that we assigned zero distance for the large deletions between host and parasitic RNAs. A point represents each genotype, and the colors of points represent the rounds of their appearance, consistent with the colors of the markers in Fig. 2A. A black line connects a pair of genotypes one Hamming distance apart in the same RNA class (round-by-round data are shown in Supplementary Figure 5). The host RNA genotypes gradually became distant from the original genotype (Host-0) as rounds proceeded (Fig. 2C and Supplementary Figure 5). From round 0 to round 43, sequences diversified around the original genotype. Then, until round 72, most of the genotypes move towards the upper-right branches. But at round 86, where a certain fraction of the genotypes shifted to the left branch and dominated until round 99. At round 104, most of the genotypes moved back to the right branch again and stayed until round 115. These frequent changes of dominant lineages imply that the fittest genotype changes frequently during the long-term replication experiment.

The population of the parasite-α represented a distinct cluster from host RNA populations (Fig.2D) and most of the genotypes are connected (i.e., one Hamming distance apart). The parasite-α did not show clear directionality throughout the long-term replication experiment (round-by-round data is shown in Supplementary Figure 5). Interestingly, we identified 18 unique mutations specific to the parasite-α (Supplementary Figure 4), not found in the corresponding region of the host sequence. The persistence of parasite-α unique mutations indicates that most of the new parasite-α genotypes were not newly generated from evolving host RNAs, but the parasite-α maintained its own lineage and evolved independently of the host RNA.

The populations of the parasite-β and -γ formed distinct clusters (Fig. 2E) and most of the genotypes were closely related (connected with one Hamming distance lines) within each class. Sequences of some parasite-β and -γ were perfectly matched with some dominant host RNAs coexisting in the same rounds as connected with broken blue lines, suggesting that these parasitic RNAs originated from these host RNAs. Unlike the parasite-α, we found only 1 and 2 unique mutations for the parasite-β and -γ (Supplementary Figure 4).

### Competitive replication assay of the host and parasitic RNAs

The diversification of host genotypes and the appearance of novel parasite classes can be a consequence of the coevolution between the hosts and parasites to adapt to each other. To test this possibility, we performed a series of competitive replication assays by using three representative host and parasitic RNAs. As the representatives, we chose the most dominant genotypes at rounds 0, 99, and 115 for host RNA (Host-0, Host-99, and Host-115, respectively). For the parasite-α, -β, and -γ, we chose the most dominant genotype for each at round 13, 99, and 115 (Parasite-α13, Parasite-β99, Parasite-γ115), respectively (sequences are available in Supplementary Text). We mixed a pair of the host and parasitic RNAs according to their order of appearance at an equivalent molar and performed competitive replication. The concentrations of replicated RNAs were measured by sequence-specific RT-qPCR (Fig. 3A). In the first pair (Host-0 vs Parasite-α13), Host-0 hardly replicated (less than 2-fold) and Parasite-α13 predominantly replicated (~200-fold), indicating that Parasite-α13 severely inhibits the original host replication, whereas in the second pair (Host-99 vs Parasite-α13), Host-99, efficiently replicated (~700-fold) with negligible replication of Parasite-α13, indicating that Host-99 acquired the resistance to Parasite-α13. In the third pair (Host-99 vs Parasite-β99), Host-99 still replicated efficiently (~1000-fold) but Parasite-β99, also replicated up to approximately 20-fold, indicating that Parasite-β99 acquired the ability to co-replicate with Host-99. In the fourth pair (Host-115 vs Parasite-β99), Host-115 repressed the replication of Parasite-β99 to less than 2-fold, indicating that Host-115 acquired the ability to evade co-replication of Parasite-β99. In the final pair (Host-115 vs Parasite-γ115), Parasite-γ115 acquired the ability to replicate up to approximately 20-fold with Host-115. These results demonstrated that successive counter-adaptive evolution (i.e., evolutionary arms-races) occurred among the host and parasitic RNAs as schematically illustrated in Fig. 3B.

**Fig.3.**
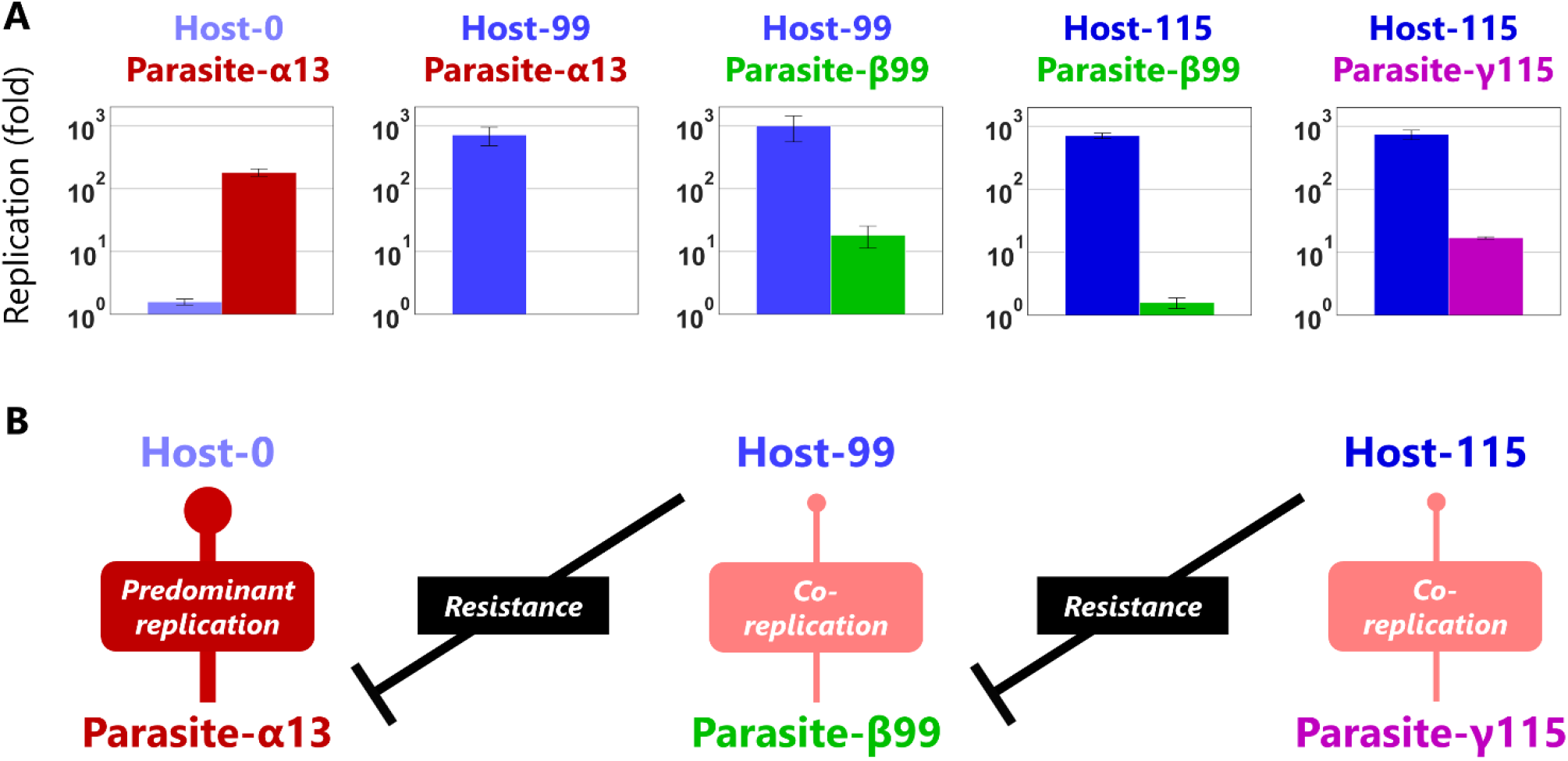
Evolutionary arms races between host and parasitic RNAs. (A) Competitive replication assay of each pair of the evolved host and parasitic RNA clones. The RNA replication reactions were performed with 10 nM of the host and parasitic RNAs for 3 h and each concentration was measured by sequence-specific RT-qPCR. Error bars represent standard errors of three independent competition assays. (B) Schematic representation of the host-parasite relationships among the RNA clones.

## Discussion

Coevolution of host and parasitic replicators is a major evolutionary driver in the evolution of life. Here, we investigated the Darwinian evolution process of an RNA replication system and demonstrated the emergence of a host-parasite ecosystem, in which new types of host and parasitic RNAs appeared one after another and exhibited antagonistic coevolutionary dynamics. During the long-term replication experiment, the host RNA continued to evolve and diversified in a sequence space. This process stands in sharp contrast to the previously reported unidirectional and rapidly slowing down evolution of the host RNA in the absence of frequent interactions with the parasitic RNAs (Ichihashi et al., 2013). The dynamic change of the host-parasite genotypes (Fig. 2C-E and Supplementary Figure 5) and phenotypes (Fig. 3) indicates that evolving parasites could have provided varying selection pressure and driven the diversification of the host RNA. The coevolution driven diversification is consistent with the consequence of natural host-parasite coevolution (Bohannan and Lenski, 2000; Buckling and Rainey, 2002; Ebert, 2008; Thompson, 1999; Woolhouse et al., 2002) and the simulated evolution in silico (Takeuchi and Hogeweg, 2008; Zaman et al., 2011). Evolutionary arms races between host-parasite molecules may be an important mechanism to generate and maintain diversity in molecular ecosystems before the origin of life.

The mutational patterns of the host and parasitic RNAs in this study posed an interesting possibility that parasites could bring about new information in a molecular population. The parasite-α accumulated 9 dominant mutations in the 3’-UTR, where the host RNA never accumulated dominant mutations through the long-term evolution (Supplementary Figure 4). This result suggests that mutations in the 3’-UTR of the host RNA are severely limited (e.g. constraint imposed by translation efficiency). Evolving and persistent parasitic molecules with less mutational constraints might add genetic novelties in the whole molecular ensemble and play a role in the evolution of complexity (Adami et al., 2000).

The evolution of genome size is another conundrum of biological complexity (Petrov, 2001; Sharov, 2006). In our study, parasites with longer genome appeared after the long-term evolution (after 94 round). Because the new parasites shared a part of the M-site sequence lacking in the parasite-α, a recognition site for Qβ replicase (Meyer et al., 1981; Schuppli et al., 1998), we can think of a plausible arms-race evolution scenario as following: (1) the parasite-α first invaded the system taking advantage of its short genome for faster replication (2) the host RNA evolved the specificity of Qβ replicase to host-specific sequences (including the M-site) to circumvent the parasite-α (3) the new parasites invaded the system because they harbored evolved M-sites recognized by evolved Qβ replicases. If this speculation is correct, the new parasites expanded the genomic information to cope with the evolved strategy of the host RNA. Recent theoretical studies suggest that host-parasite antagonistic coevolution is an effective mechanism to increase the complexity of individuals (Luis and Ricard, 2019; Zaman et al., 2014). Whether the host RNA expands the genome to avoid the new parasites remains an interesting question.

A typical phenomenon in host-parasite coevolution is Red Queen dynamics (Rabajante et al., 2015; van Valen, 2007), in which host and parasite populations oscillate due to persistent replacement of dominant hosts and parasites. The host-parasite RNA population in our replication experiment exhibited Red Queen dynamics with a remarkable feature of damping fluctuations. One possibility of the damped oscillation is the elevation of the average parasite resistance in the evolved host RNA population against the parasitic RNAs, which might be partly supported by the weakened inhibition of the host replication by the parasitic RNAs in later rounds (Fig. 3). Another possibility is that the increased diversity (Fig. 2C-E, Supplementary Figure 5) allows competition among various types of host and parasitic RNAs to average the population dynamics. A study on *Daphnia* and its parasite also reported damped long-term host-parasite Red Queen coevolutionary dynamics and suggested that the increased host diversity as a consequence of coevolution could decrease fluctuations in host-parasite Red Queen dynamics (Decaestecker et al., 2013). Our simple and fast-evolving host-parasite RNA replication system may offer a useful platform to investigate the relationship between ecological and evolutionary dynamics of hosts and parasites and further pursue an exciting evolution scenario, such as the emergence of cooperation between host-parasite replicators (Ikegami and Kaneko, 1990).

## Materials and Methods

### A long-term replication experiment

In this study, we additionally performed 77 rounds of replication using the RNA population at round 43 of the previous experiment by the same method (Bansho et al., 2016). In this method, initially, 10 μL of the reconstituted *E. coli* translation system (Shimizu et al., 2001) containing 1 nM of the original host RNA (Host-0, the round 128 clone in a previous study (Ichihashi et al., 2013) was mixed with 1 mL of the buffer-saturated oil prepared as described previously (Ichihashi et al., 2013) using a homogenizer (POLYTRON PT-1300D; KINEMATICA) at 16,000 rpm for 1 min on ice. The water-in-oil droplet was incubated at 37 °C for 5 h to induce the protein translation reaction and the RNA replication reaction. For the next round of RNA replication, a fraction of the water-in-oil droplets (200 μL) was transferred and mixed with the new buffer-saturated oil (800 mL) and the new translation system (10 μL) using the homogenizer at 16,000 rpm for 1 min on ice, then incubated at 37 °C for 5 h. The average diameter of the water-in-oil droplets was approximately 2 μm (Bansho et al., 2016). After the incubation step in each round, RNA concentrations were measured as described below. The composition of the translation system is described in the previous study (Bansho et al., 2016).

### Measurement of host RNA concentrations

The water-in-oil droplets after the incubation step were diluted 10000-fold with 1 mM EDTA (pH 8.0) and subjected to RT-qPCR (PrimeScript One Step RT-PCR Kit (TaKaRa)) with primer 1 and 2 after heating at 95 °C for 5 min. These primers specifically bind to the host RNA. To draw a standard curve in RT-qPCR, dilution series of the water-in-oil droplets containing the original host RNA diluted 10000-fold with 1 mM EDTA were used.

### Measurement of parasitic RNA concentrations

To determine the concentrations of the parasitic RNAs that appeared during the long-term replication experiment (Fig. 2A), polyacrylamide gel electrophoresis was performed followed by quantification of the fluorescent intensities of the parasitic RNA bands using ImageJ. The water phases were collected from the water-in-oil droplets after the incubation step at each round and RNAs were purified with spin columns (RNeasy, QIAGEN). The purified RNA samples and dilution series of the standard parasitic RNA (S222 RNA (Hosoda et al., 2007)) were subjected to 8% polyacrylamide gel electrophoresis with 0.1% SDS in TBE buffer (pH 8.4) containing Tris(hydroxymethyl)aminomethane (100 mM), boric acid (90 mM), and EDTA (1 mM), followed by staining with SYBR Green II (Takara). The fluorescent intensities of the parasitic RNA bands were quantified and the concentrations were determined based on the standard curve drawn with the dilution series of the standard parasitic RNA bands.

In the previous study (Bansho et al., 2016), we measured the parasitic RNA concentration from the replication kinetics using a purified Qβ replicase and the detection limit was lower than that of this study. This method could not be employed here because it is unable to distinguish different classes of parasitic RNAs appeared in this study.

### Sequence analysis

The RNA mixture of round-13, 24, 33, 39, 43, 50, 53, 60, 65, 72, 86, 91, 94, 99, 104, 110, and 115 in the long-term replication experiment were purified with spin columns (RNeasy, QIAGEN). The purified RNAs were reverse-transcribed using PrimeScript reverse transcriptase (Takara) and primer 3 and then PCR-amplified using primer 3 and 4. The PCR products were subjected to agarose gel electrophoresis and the bands corresponding to the host cDNA and the parasitic cDNA were separately extracted by using E-gel CloneWell (Thermo Fisher Scientific). The host and the parasite-β and -γ were sequenced using PacBio RS II with C4/P6 chemistry (Pacific Biosciences), and the parasite-α was sequenced using MiSeq (Illumina). To reduce read errors in the PacBio RS II sequencing, we used circular consensus sequencing (CCS) reads at least comprising five and ten reads for the host and the parasites, respectively. All the sequence reads were subjected to sequence alignment with a reference sequence (the original host sequence) separately for each molecular species (i.e. the host, the parasite-α, -β, and -γ) using MAFFT v7.294b with FFT-NS-2 algorithm (K Katoh, K Misawa, K Kuma, 2002). Frequencies of mutations were calculated for each round sample and 74 dominant mutations were identified (Supplementary Figure 4). These 74 dominant mutations are located in 72 sites (i.e., a few mutations were introduced in the same sites). In the following analysis, we focused on only the genotypes consist of these 72 sites with 74 mutations.

### Mapping dominant genotypes on a two-dimensional space

From the genotypes consisting of the 72 sites with the 74 mutations, the top-90 most dominant genotypes were identified for each host and parasitic species at each round. Hamming distances between all the pairs of the genotypes were calculated and a square distance matrix *D*, whose *i,j*-th component *d_ij_* represents the square of the Hamming distance between the *i*-th genotype and the *j*-th genotype, was constructed. Using principal coordinate analysis on the square distance matrix *D*, the positions of each genotype were determined. The matrix *D* was transformed into a kernel matrix *K* = −1/2*CDC*, where *C* is the centering matrix. Let *λ_k_* and *e_k_* ≡ (*e_k1_*, *e_k2_*,…,*e_kM_*) denote the *k*-th eigenvalue and the *k*-th normalized eigenvector, where *λ_1_* > *λ_2_* >…> *λ_M_* and | *e_k_* | = 1 for all *k* and *M* is the dimension of the kernel matrix *K*. The eigenvalues and eigenvectors of the kernel matrix *K* were calculated and the *i*-th genotype was plotted in the two-dimensional

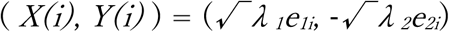

space with a coordinate described as follows:

### Competitive replication assay of host and parasitic RNAs

The six plasmids, each containing T7 promoter and following cDNA sequences of Host-0, Host-99, Host-115, Parasite-α13, Parasite-β99, and Parasite-γ115, were constructed using the gene synthesis service of Eurofins Genomics. Each RNA was synthesized by *in vitro* transcription with T7 RNA polymerase (TaKaRa) from the plasmids digested with SmaI according to a previous study (Yumura et al., 2017). We mixed 10 nM of each host and parasitic RNAs in the cell-free translation system and incubated 37 °C for 3 h. The concentrations of the host and the parasitic RNAs were measured by RT-qPCR (PrimeScript One Step RT-PCR Kit (TaKaRa)) with sequence-specific primers (Supplementary text).

## Supporting information

Genotype frequency data

## Acknowledgements

We would like to thank Nobuto Takeuchi, Kunihiko Kaneko, Ryo Mizuuchi, Yoshihiro Sakatani, Yannick Rondelez, and Tetsuya Yomo for useful discussions and comments. This work was supported by JSPS KAKENHI grant numbers JP15KT0080, JP15H04407 and JP17J01023.

## Author contributions

T.F. and N.I. designed the research. T.F., K.U., Y.B., D.M., S.N. conducted the experiments. T.F. analyzed the data. T.F and N.I. wrote the manuscript.

## References

Adami C, Ofria C, Collier TC. 2000. Evolution of biological complexity. Proceedings of the National Academy of Sciences of the United States of America. doi:10.1073/pnas.97.9.4463

Bansho Y, Furubayashi T, Ichihashi N, Yomo T. 2016. Host-parasite oscillation dynamics and evolution in a compartmentalized RNA replication system. Proceedings of the National Academy of Sciences of the United States of America 113. doi:10.1073/pnas.1524404113

Bansho Y, Ichihashi N, Kazuta Y, Matsuura T, Suzuki H, Yomo T. 2012. Importance of parasite RNA species repression for prolonged translation-coupled RNA self-replication. Chemistry and Biology. doi:10.1016/j.chembiol.2012.01.019

Bergh Ø, Børsheim KY, Bratbak G, Heldal M. 1989. High abundance of viruses found in aquatic environments. Nature. doi:10.1038/340467a0

Bohannan BJM, Lenski RE. 2000. Linking genetic change to community evolution: Insights from studies of bacteria and bacteriophage. Ecology Letters. doi:10.1046/j.1461-0248.2000.00161.x

Bresch C, Niesert U, Harnasch D. 1980. Hypercycles, parasites and packages. Journal of Theoretical Biology 85:399–405. doi:10.1016/0022-5193(80)90314-8

Buckling A, Rainey PB. 2002. Antagonistic coevolution between a bacterium and a bacteriophage. Proceedings of the Royal Society B: Biological Sciences. doi:10.1098/rspb.2001.1945

Chetverin AB, Chetverina H v, Demidenko AA, Ugarov VI. 1997. Nonhomologous RNA recombination in a cell-free system: evidence for a transesterification mechanism guided by secondary structure. Cell 88:503–513. doi:10.1016/s0092-8674(00)81890-5

Claverie JM. 2006. Viruses take center stage in cellular evolution. Genome Biology. doi:10.1186/gb-2006-7-6-110

Decaestecker E, de Gersem H, Michalakis Y, Raeymaekers JAM. 2013. Damped long-term host-parasite Red Queen coevolutionary dynamics: A reflection of dilution effects? Ecology Letters. doi:10.1111/ele.12186

Deininger PL, Moran J v, Batzer MA, Kazazian HH. 2003. Mobile elements and mammalian genome evolution. Current opinion in genetics & development 13:651–658.

Ebert D. 2008. Host-parasite coevolution: Insights from the Daphnia-parasite model system. Current Opinion in Microbiology. doi:10.1016/j.mib.2008.05.012

Elbarbary RA, Lucas BA, Maquat LE. 2016. Retrotransposons as regulators of gene expression. Science 351:aac7247–aac7247. doi:10.1126/science.aac7247

Forterre P, Prangishvili D. 2013. The major role of viruses in cellular evolution: facts and hypotheses. Current opinion in virology 3:558–565. doi:10.1016/j.coviro.2013.06.013

Forterre P, Prangishvili D. 2009. The great billion-year war between ribosome- and capsid encoding organisms (cells and viruses) as the major source of evolutionary noveltiesAnnals of the New York Academy of Sciences. doi:10.1111/j.1749-6632.2009.04993.x

Furubayashi T, Ichihashi N. 2018. Sustainability of a compartmentalized host-parasite replicator system under periodic washout-mixing cycles. Life 8. doi:10.3390/life8010003

García-Villada L, Drake JW. 2012. The Three Faces of Riboviral Spontaneous Mutation: Spectrum, Mode of Genome Replication, and Mutation Rate. PLoS Genetics 8:e1002832. doi:10.1371/journal.pgen.1002832

Higgs PG, Lehman N. 2015. The RNA World: Molecular cooperation at the origins of life. Nature Reviews Genetics. doi:10.1038/nrg3841

Hosoda K, Matsuura T, Kita H, Ichihashi N, Tsukada K, Yomo T. 2007. Kinetic analysis of the entire RNA amplification process by Qβ replicase. Journal of Biological Chemistry 282. doi:10.1074/jbc.M700307200

Ichihashi N, Usui K, Kazuta Y, Sunami T, Matsuura T, Yomo T. 2013. Darwinian evolution in a translation-coupled RNA replication system within a cell-like compartment. Nature Communications. doi:10.1038/ncomms3494

Ikegami T, Kaneko K. 1990. Computer symbiosis-emergence of symbiotic behavior through evolution. Physica D: Nonlinear Phenomena 42:235–243. doi:10.1016/0167-2789(90)90077-3

Iranzo J, Lobkovsky AE, Wolf YI, Koonin E v. 2014. Virus-host arms race at the joint origin of multicellularity and programmed cell death. Cell cycle (Georgetown, Tex) 13:3083–3088. doi:10.4161/15384101.2014.949496

Joyce GF, Szostak JW. 2018. Protocells and RNA self-replication. Cold Spring Harbor Perspectives in Biology. doi:10.1101/cshperspect.a034801

K Katoh, K Misawa, K Kuma TM. 2002. MAFFT: a novel method for rapid multiple sequence alignment based on fast Fourier transform. Nucleic acids research.

Koonin E v, Dolja V v. 2013. A virocentric perspective on the evolution of life. Current opinion in virology 3:546–557. doi:10.1016/j.coviro.2013.06.008

Koonin E v, Wolf YI, Katsnelson MI. 2017. Inevitability of the emergence and persistence of genetic parasites caused by evolutionary instability of parasite-free states. Biology direct 12:31. doi:10.1186/s13062-017-0202-5

Koskella B, Brockhurst MA. 2014. Bacteria-phage coevolution as a driver of ecological and evolutionary processes in microbial communities. FEMS Microbiology Reviews 38:916–931. doi:10.1111/1574-6976.12072

Luis FS, Ricard VS. 2019. How parasites expand the computational landscape of life. arXiv.

Matsumura S, Kun A, Ryckelynck M, Coldren F, Szilagyi A, Jossinet F, Rick C, Nghe P, Szathmary E, Griffiths AD. 2016. Transient compartmentalization of RNA replicators prevents extinction due to parasites. Science 354:1293–1296. doi:10.1126/science.aag1582

Meyer F, Weber H, Weissmann C. 1981. Interactions of Qβ replicase with Qβ RNA. Journal of Molecular Biology 153:631–660. doi:10.1016/0022-2836(81)90411-3

Mills DR, Kramer FR, Spiegelman S. 1973. Complete nucleotide sequence of a replicating RNA molecule. Science. doi:10.1126/science.180.4089.916

Mizuuchi R, Ichihashi N. 2018. Sustainable replication and coevolution of cooperative RNAs in an artificial cell-like system. Nature Ecology and Evolution. doi:10.1038/s41559-018-0650-z

Müller V, de Boer RJ, Bonhoeffer S, Szathmáry E. 2018. An evolutionary perspective on the systems of adaptive immunity. Biological Reviews. doi:10.1111/brv.12355

Orgel LE. 1992. Molecular replication. Nature 358:203–209. doi:10.1038/358203a0

Petrov DA. 2001. Evolution of genome size: New approaches to an old problem. Trends in Genetics. doi:10.1016/S0168-9525(00)02157-0

Rabajante JF, Tubay JM, Uehara T, Morita S, Ebert D, Yoshimura J. 2015. Red Queen dynamics in multi-host and multi-parasite interaction system. Scientific Reports. doi:10.1038/srep10004

Schuppli D, Miranda G, Qiu S, Weber H. 1998. A branched stem-loop structure in the M-site of bacteriophage Qβ RNA is important for template recognition by Qβ replicase holoenzyme. Journal of Molecular Biology 283:585–593. doi:10.1006/jmbi.1998.2123

Sharov AA. 2006. Genome increase as a clock for the origin and evolution of life. Biology Direct. doi:10.1186/1745-6150-1-17

Shimizu Y, Inoue A, Tomari Y, Suzuki T, Yokogawa T, Nishikawa K, Ueda T. 2001. Cell-free translation reconstituted with purified components. Nature Biotechnology 19:751–755. doi:10.1038/90802

Suttle CA. 2007. Marine viruses — major players in the global ecosystem. Nature Reviews Microbiology 5:801–812. doi:10.1038/nrmicro1750

Szathmáry E, Demeter L. 1987. Group selection of early replicators and the origin of life. Journal of theoretical biology 128:463–486.

Szathmáry E, Maynard Smith JM. 1997. From replicators to reproducers: The first major transitions leading to life. Journal of Theoretical Biology. doi:10.1006/jtbi.1996.0389

Takeuchi N, Hogeweg P. 2009. Multilevel Selection in Models of Prebiotic Evolution II: A Direct Comparison of Compartmentalization and Spatial Self-Organization. PLoS Computational Biology 5:e1000542. doi:10.1371/journal.pcbi.1000542

Takeuchi N, Hogeweg P. 2008. Evolution of complexity in RNA-like replicator systems. Biology Direct. doi:10.1186/1745-6150-3-11

Thompson JN. 1999. Specific hypotheses on the geographic mosaic of coevolution. American Naturalist. doi:10.1086/303208

van Valen L. 2007. New Evolutionary low.

Wochner A, Attwater J, Coulson A, Holliger P. 2011. Ribozyme-catalyzed transcription of an active ribozyme. Science. doi:10.1126/science.1200752

Woolhouse MEJ, Webster JP, Domingo E, Charlesworth B, Levin BR. 2002. Biological and biomedical implications of the co-evolution of pathogens and their hosts. Nature Genetics. doi:10.1038/ng1202-569

Yumura M, Yamamoto N, Yokoyama K, Mori H, Yomo T, Ichihashi N. 2017. Combinatorial selection for replicable RNA by Qβ replicase while maintaining encoded gene function. PLoS ONE. doi:10.1371/journal.pone.0174130

Zaman L, Devangam S, Ofria C. 2011. Rapid host-parasite coevolution drives the production and maintenance of diversity in digital organismsGenetic and Evolutionary Computation Conference, GECCO’11. doi:10.1145/2001576.2001607

Zaman L, Meyer JR, Devangam S, Bryson DM, Lenski RE, Ofria C. 2014. Coevolution Drives the Emergence of Complex Traits and Promotes Evolvability. PLoS Biology. doi:10.1371/journal.pbio.1002023

