## Supplementary information for "Emergence and diversification of a host-parasite RNA ecosystem through Darwinian evolution"

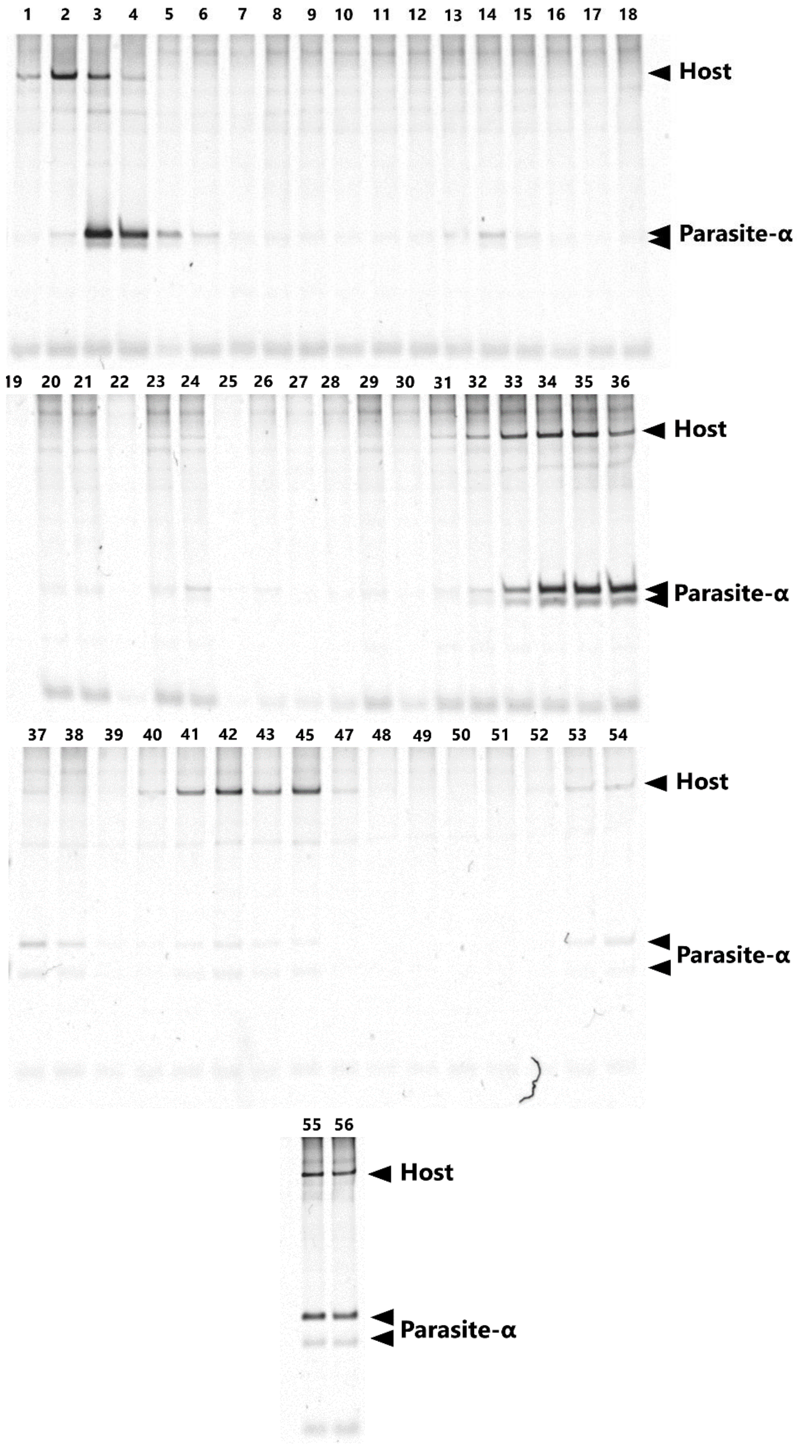


**Supplementary Figure 1. The native polyacrylamide gel electrophoresis of the RNA mixture during the long-term replication experiment.**

The numbers above the gels indicate the sampled rounds. The parasitic RNAs exhibit multiple bands due to structural heterogeneity.

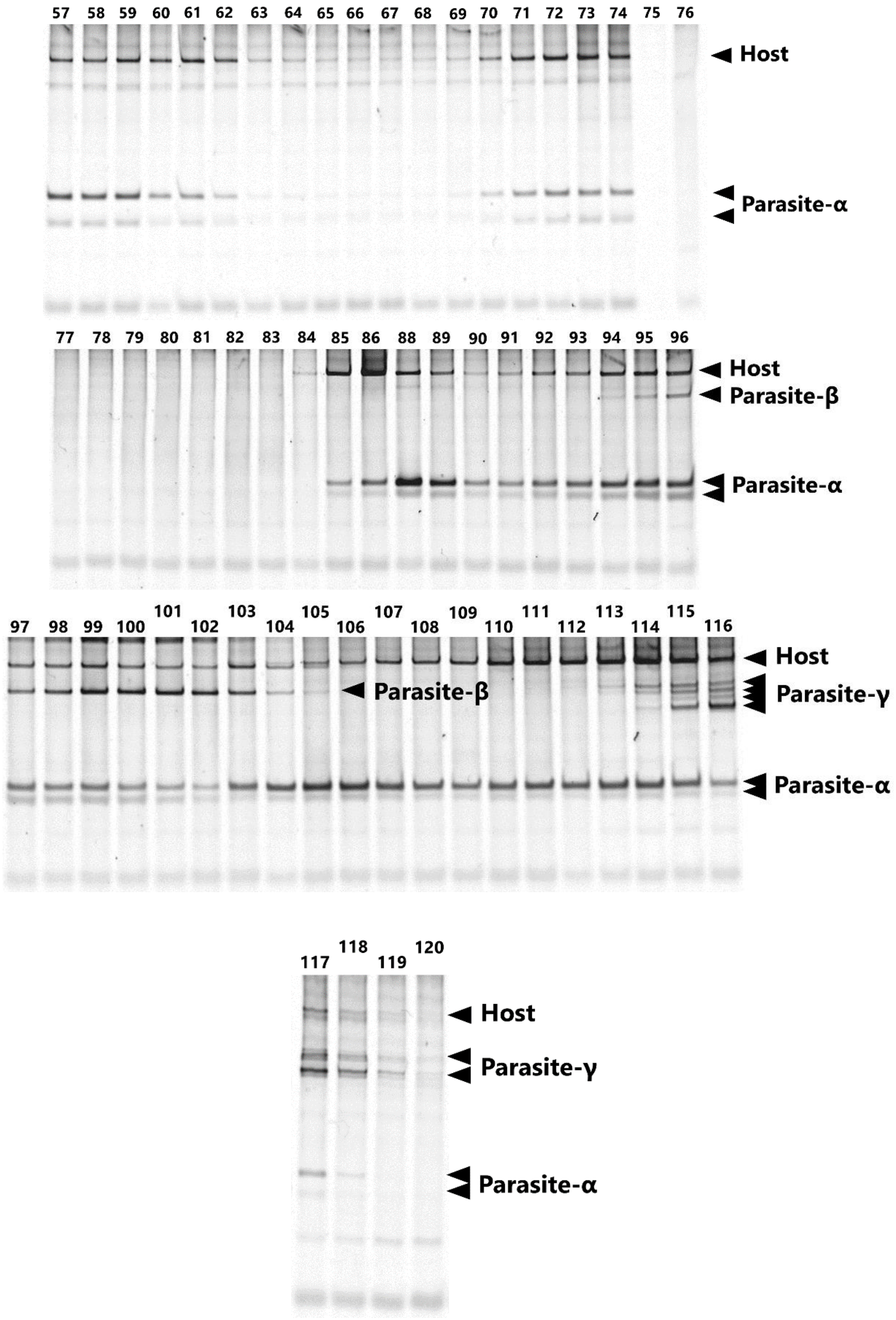


**Supplementary Figure 1 (continued).**

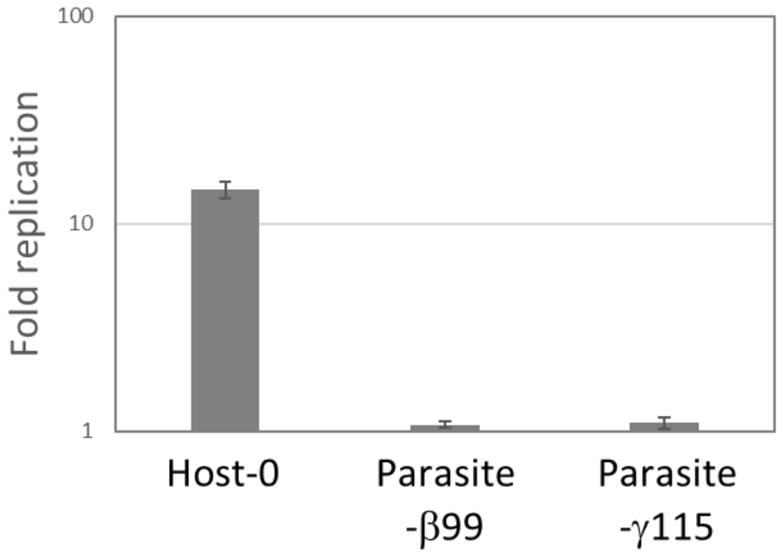


**Supplementary Figure 2. Replication of Parasite-β99 and Parasite-γ115 without host species.**
The RNA replication reactions were performed with 10 nM of each RNA for 5 h, and RNA concentration was measured by sequence-specific RT-qPCR. Error bars represent standard errors of three independent competition assays. Host-0 was also replicated for comparison.

**Host**

(PacBio RS Ⅱ)

| Round | Read number |
|---|---|
| 13 | 4143 |
| 24 | 605 |
| 33 | 718 |
| 39 | 365 |
| 43 | 484 |
| 50 | 1020 |
| 53 | 1358 |
| 60 | 1409 |
| 65 | 3097 |
| 72 | 1364 |
| 86 | 2135 |
| 91 | 637 |
| 94 | 2058 |
| 99 | 855 |
| 104 | 2535 |
| 110 | 1758 |
| 115 | 4003 |

**Parasite-α**

(MiSeq)

| Round | Read number |
|---|---|
| 13 | 5000 |
| 24 | 4990 |
| 33 | 4999 |
| 43 | 4986 |
| 53 | 4987 |
| 60 | 4963 |
| 72 | 4994 |
| 86 | 4974 |
| 91 | 4998 |
| 94 | 4995 |
| 99 | 4984 |
| 104 | 4926 |
| 110 | 4929 |
| 115 | 4959 |

**Parasite-β**

(PacBio RS Ⅱ)

| Round | Read number |
|---|---|
| 99 | 4999 |

**Parasite-γ**

(PacBio RS Ⅱ)

| Round | Read number |
|---|---|
| 115 | 1753 |

**Supplementary Figure 3. Read numbers of deep sequencing.**
Note that the coverage is 100% for all the reads.

| Mutation index | Base change | Amino acid change (host) | Appearance in different classes |
|---|---|---|---|
| 1 | A40G | | α |
| 2-1<br>2-2 | G46A<br>G46deletion | | α<br>Host |
| 3 | A49G | | Host, β |
| 4 | A53G | | Host, β |
| 5 | G55deletion | | Host, α, β, γ |
| 6 | C67U | | α |
| 7 | C72U | | Host, α, γ |
| 8 | U78A | | Host, α, β |
| 9 | C86U | | Host, α, β, γ |
| 10 | U114C | | Host, α, β, γ |
| 11 | C115A | | α |
| 12 | A116G | | Host, α, β |
| 13 | insertion129A | | Host, α, β, γ |
| 14 | G141A | | Host, α, β, γ |
| 15 | C152deletion | | α |
| 16 | U156C | | α |
| 17 | C158A | | α |
| 18 | U159deletion | | α |
| 19 | C160U<br>C160A | | α |
| 20 | U161C | | Host, β |
| 21 | G172U | | Host |
| 22 | G178C | | γ |
| 23 | C188U | | Host, β |
| 24 | U198C | | Host |
| 25 | U205C | | Host |
| 26 | A215G | | Host |
| 27 | A225G | | Host, β |
| 28 | C232U | Pro2Ser | Host |
| 29 | C259U | Leu11Phe | Host, β |
| 30 | A294G | synonymous | Host |
| 31 | insertion296U | frameshift | Host |
| 32 | U321deletion | frameshift | Host |
| 33 | C399U | Pro57Leu | Host |
| 34 | C414deletion | frameshift | Host, β |
| 35 | A438G | Tyr70Cys | Host, β |
| 36 | A452G | Ile75Val | Host, β |
| 37 | U501C | Val91Ala | Host |
| 38 | G565A | synonymous | Host, β, γ |
| 39 | U568C | synonymous | Host |
| 40 | G599U | Gly124Cys | Host |
| 41-1<br>41-2 | A626G<br>A626C | Arg133Gly<br>synonymous | Host, β<br>Host, γ |
| 42 | A631G | synonymous | Host |
| 43 | U672C | Met148Thr | Host |
| 44 | U683C | Cys152Arg | β |
| 45 | U692C | Ser155Pro | Host, γ |
| 46 | A706G | synonymous | Host |
| 47 | A751G | synonymous | Host |
| 48 | G761A | Ala178Thr | β |
| 49 | A851G | Lys208Glu | Host |
| 50 | U920deletion | frameshift | Host |
| 51 | C1103deletion | frameshift | Host |
| 52 | U1116C | Phe296Ser | Host |
| 53 | U1117C | synonymous | Host |
| 54 | U1190C | Ser321Pro | Host |
| 55 | C1246A | synonymous | Host |
| 56 | U1280C | Ser351Pro | Host |
| 57 | G1399deletion | frameshift | Host |
| 58 | U1568C | Tyr447His | Host |
| 59 | U1572G | Leu448Arg | Host |
| 60 | A1605G | Gln459Arg | Host |
| 61 | A1608G | His460Arg | Host |
| 62 | U1633C | synonymous | Host |
| 63 | U1720A | His497Gln | Host |
| 64 | C1982A | | α |
| 65 | A1983G | | α |
| 66 | insertion1985U | | α |
| 67 | C1986U | | α |
| 68 | A1987G | | α |
| 69 | G1989deletion | | α |
| 70 | G2001C | | α |
| 71 | insertion2002U | | α |
| 72 | U2010C | | α |

**Supplementary Figure 4. List of dominant mutations.**

The untranslated regions and the coding region are indicated with red and blue backgrounds respectively. The base numbers are determined based on the single reference sequence (shown in the Supplemental text), which contains all dominant insertions (129A, 296U, 1985T, and 2002T) in the original host sequence (Host-0). α, β, and γ in the "Appearance in different species" are shorthand expressions for the parasite-α, -β, and -γ.

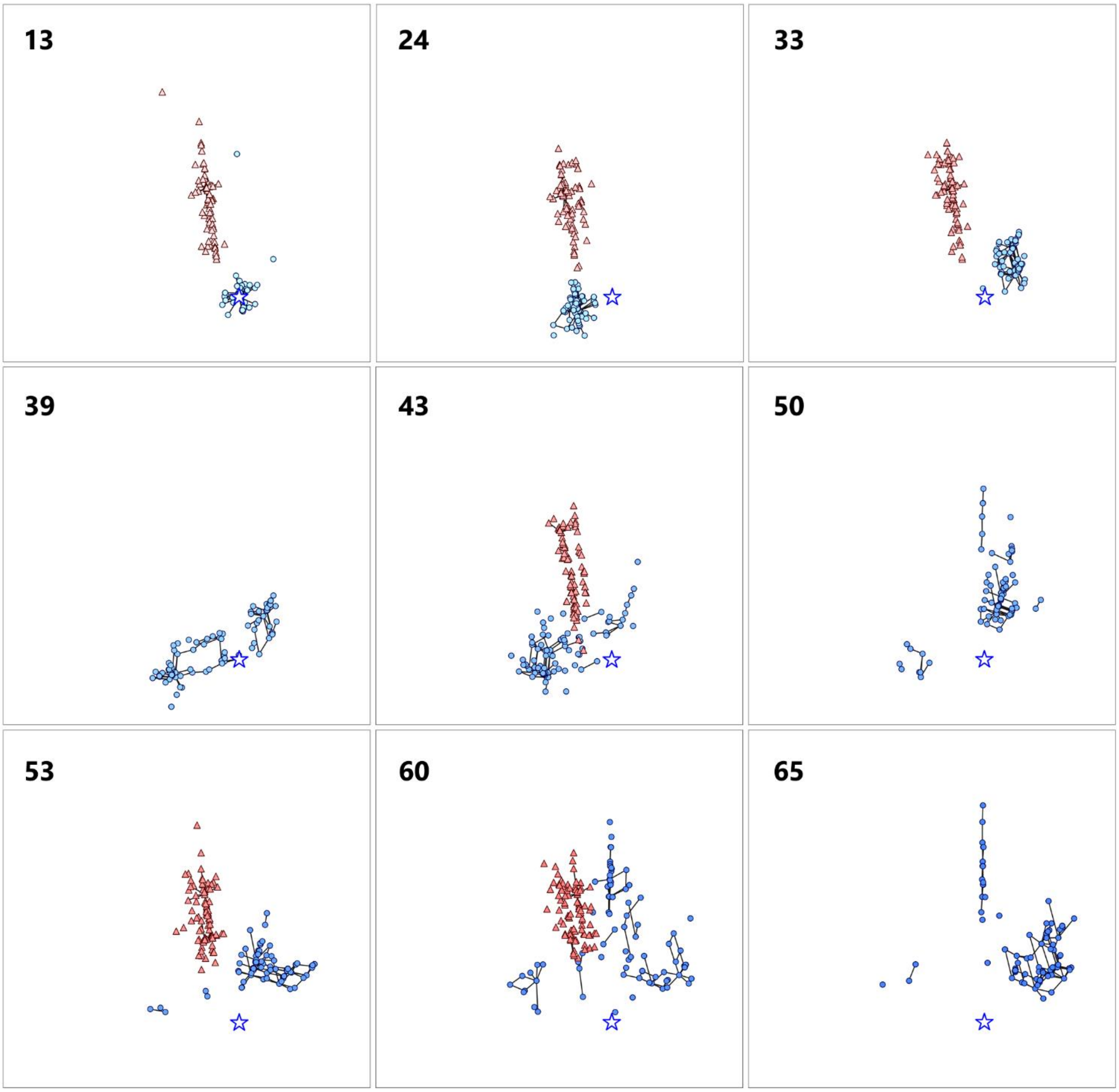


**Supplementary Figure 5. Round-by-round top-90 host and parasite-α genotypes on the 2D space.** A star in each figure represents the coordinate of the original host genotype (Host-0). Black lines connect pairs of points one Hamming distance apart in the same RNA species. A broken blue line connects a pair of points zero Hamming distance apart in the different RNA species, ignoring the large deletion of the parasite-α. Parasite-α is not shown in the round-39, 50, and 65 because they could not have been recovered and sequenced.

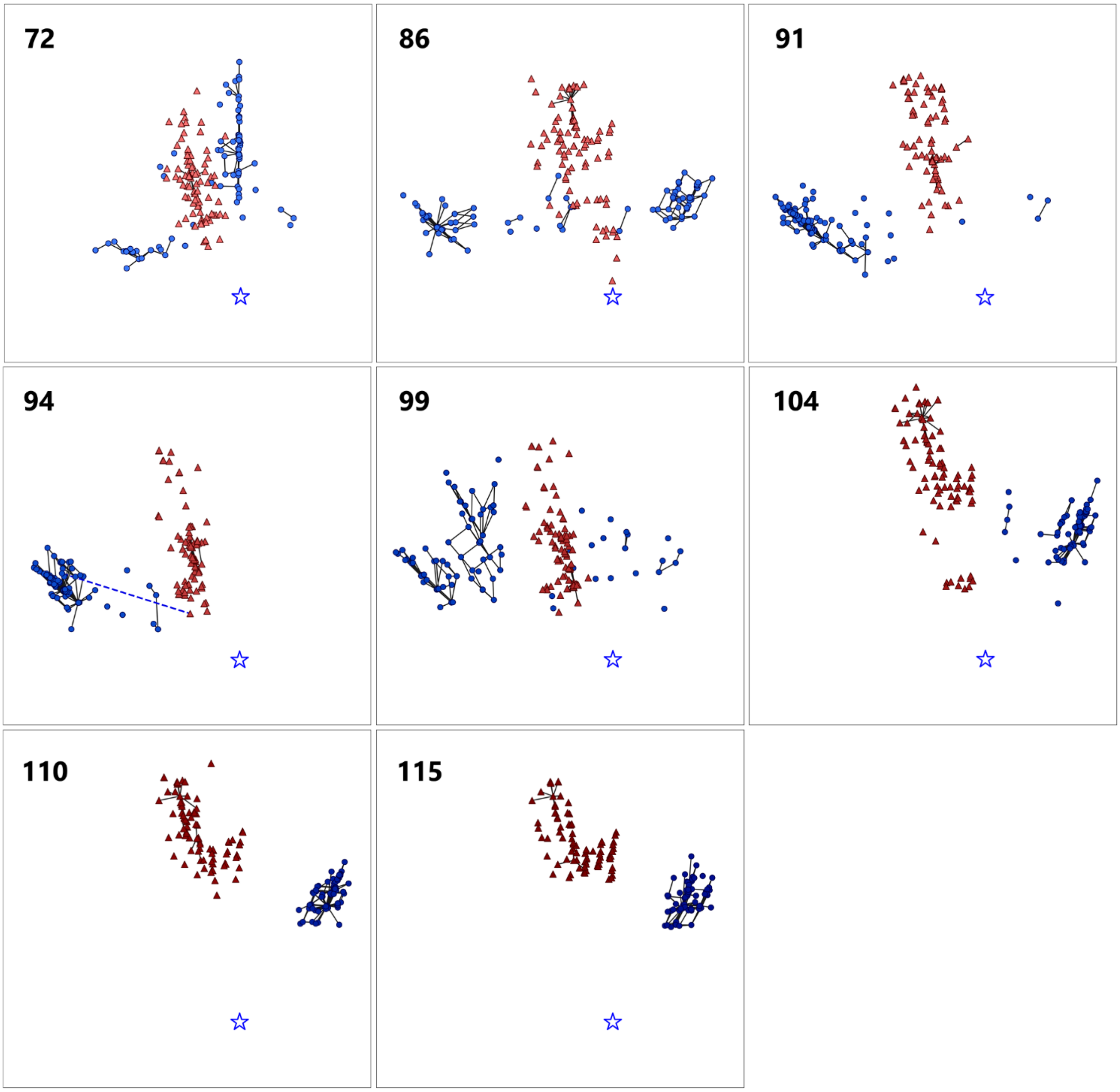


**Supplementary Figure 5 (continued).**

| Mutation index | Host-99 | Host-115 | Amino acid change |
|---|---|---|---|
| 3 | + | − | 5'-UTR |
| 4 | + | − | |
| 7 | − | + | |
| 8 | + | − | |
| 9 | + | + | |
| 10 | + | + | |
| 12 | + | − | |
| 13 | − | + | |
| 14 | − | + | |
| 27 | + | − | |
| 29 | − | + | Leu11Phe |
| 37 | − | + | Val91Ala |
| 38 | − | + | synonymous |
| 40 | + | − | Gly124Cys |
| 41 | − | + | synonymous |
| 42 | + | − | synonymous |
| 47 | + | − | synonymous |
| 49 | + | + | Lys208Glu |
| 56 | + | − | Ser351Pro |
| 58 | − | + | Tyr447His |
| 59 | + | − | Leu448Arg |
| 60 | + | − | Gln459Arg |
| 61 | + | − | His460Arg |

**Supplementary Figure 6. Dominant mutations in Host-99 and Host-115.**
Mutation indexes correspond to those in Supplementary Figure 4.

### Supplementary Text

**The reference sequence used to determine the position of bases in Supplementary Figure 4.**

The sequence consists of the original host sequence with four dominant insertion cites (2045 bases in total). The potential insertion positions are represented with asterisks.

GGGAACCCCCCUUCGGGGGGUCACCUCGCGCAGCGGGCUACGCGAGGGAGCCACGCUGCGAAG
CAGCGUGGCGGUUCUCGUGCGUCACCGAAACGCACGAAGGUCGCGCCUCUUCACGAGGCGUC
ACC*UGGGAGAGCGCGAAAGCGCUAGCCCGUGCUCUAGCUCUAGAAGGUCUCGAGAUCUCCUC
UAGAGAUAAUUUUGUUUAACUCUAAGAAGGAGAUAUACACAUGCCUAAGACAGCAUCUUCGC
GUAACUCUCUCAGCGCACAAUUGCGCCGAGCCGCGAACACAAGAA*UUGAGGCUGAAGGUAA
CCUCGCACUUUCCAUUGCCAACGAUUUACUGUUGGCCUAUGGUCAGUCGCCAUUUAACUCUG
AGGCUGAGUGUAUUUCAUUCAGCCCGAGAUUCGACGGGACCCCGGAUGACUUUAGGAUAAAU
UAUCUUAAAGCCGAGAUCAUGUCGAAGUAUGACGACUUCAGCCUAGGUAUUGAUACCGAAGC
UGUUGCCUGGGAGAAGUUCCUGGCAGCAGAGGCUGAAUGUGCUUUAACGAACGCUCGUCUCU
AUAGGCCUGACUACAGUGAGGAUUUCAAUUUCUCACUGGGCGAGUCAUGUAUACACAUGGCU
CGUAGAAAAAUAGCCAAGCUAAUAGGAGAUGUUCCGUCCGUUGAGGAUAUGUUGCGUCACUG
CCGAUUUUCUGGCGGUGCUACAACAACGAAUAACCGUUCGUACAGUCAUCCGUCCUUCAAGU
UUGCACUUCCGCAAGCGUGUACGCCUCGGGCUUUGAAGUAUGUUUUAGCUCUCAGAGCUUCU
ACACAUUUCGAUAUCAGAAUUUCUGAUAUUAGCCCUUUUAAUAAAGCAGUUACCGUACCUAA
GAACAGUAAGACAGAUCGUUGUAUUGCUAUCGAACCUGGUUGGAAUAUGUUUUUCCAACUGG
GUAUCGGUGGCAUUCUACGCGAUCGGUUGCGUUGCUGGGGUAUCGAUCUGAAUGAUCAGACG
AUAAAUCAGCGCCGCGCUCACGAAGGCUCCGUUACUAAUAACUUAGCAACGGUUGAUCUCUC
AGCGGCAAGCGAUUCUAUGUCUCUUGCCCUCUGUGAGCUCUUAUUGCCCCGAGGCUGGUUUG
AGGUUCUUAUGGACCUCAGAUCACCUAAGGGGCGAUUGCCUGACGGUAGUGUUGUUACCUAC
GAGAAGAUUUCUUCUAUGGGUAACGGUUACACAUUCGAGCUCGAGUCGCUUAUUUUGCUUC
UCUCGCUCGUUCCGUUUGUGAGAUACUGGACUUAGACUCGUCUGAGGUCACUGUUUACGGAG
ACGAUAUUAUUUUACCGUCCCGUGCAGUCCCUGCCCUCCGGGAAGUUUUUAAGUAUGUUGGU
UUUACGACCAAUACUAAAAAGACUUUUUCCGAGGGGCCGUUCAGAGAGUCGUGCGGCAAGCA
CUACUAUUCUGGCGUAGAUGUUACUCCCUUUUACAUACGUCACCGUAUAGUGAGUCCUGCCG
AUUUAAUACUGGUUUUGAAUAACCUAUAUCGGUGGGCCACUAUUGACGGCGUAUGGGAUCCU
AGGGCCCAUUCUGUGUACCUCAAGUAUCGUAAGUUGCUGCCUAAACAGCUGCAACAUAAUAC
UAUACCUGACGGUUACGGUGAUGGUGCCCUCGUCGGAUCGGUCCUAAUCAAUCCUUUCGCGA
AAAACCGCGGGUGGAUCCGGUACGUACCGGUGAUUACGGACCAUACAAGGGACCAAGAGCGC
GCUGAGUUGGGGUCGUAUCUCUACGACCUCUUCUCGCGUUGUCUCUCGGAAAGUAACGAUGG
GUUGCCUCUUAGGGGUCCAUCGGGUUGCGAUUUUGCUGAUCUAUUUGCCAUCGAUCAGCUUA
UCUGUAGGAGUGAUCCUACGAAGAUAAGCAGGCCUACCGGUAAAUUCGAUAUACAGUACAUC
GCGUGCUGUUGUUCGGGUUGUUGUUAGGCUUGCGGCCGCACUCGAGAGAUCUAGAGCAU*CA
CGGUCGAACUCCCG*UACGAGGUGCCCGCACCUCGUCCCCCCUUCCGGGGGGUCCCC

### Primer list and sequences

**Primers used for RT-qPCR for measurement of host RNA concentrations in the long-term replication experiment**

Primer 1 : GCTGCCTAAACAGCTGCAAC

Primer 2 : CGCTCTTGGTCCCTTGTATG

**Primers for reverse transcription and PCR to recover cDNAs for sequence analysis**

Primer 3 : CCGGAAGGGGGGACGAGG

Primer 4 : GGGTCACCTCGCGCAGC

**Primers for RT-qPCR for measurement of RNA concentrations in the competitive replication assay**
Host-0 forward : ATACACATGGCTCGTAGAAAA
Host-0 reverse : GGCGTACACGCTTGCGGAAGT
Host-99 forward : TTTGTGAGATACTGGACTTAGACC
Host-99 reverse : GCAGCAACTTACGATACTTGC
Host-115 forward : CCTAGGGCCCATTCTGTGC
Host-115 reverse : GGTTTTTCGCGAAAGGATTGA
Parasite-α13 forward : TACCGAAACGCACGAAGG
Parasite-α13 reverse : CGTACGGGAGTTCGACCG
Parasite-β99 forward : GATAAATTGTCTTAAAGCCGAGG
Parasite-β99 reverse : CTCCTATTAGCTTGGCTATTTTTCC
Parasite-γ115 forward : TAAGAAGGAGATATGCTTTAACGAA
Parasite-γ115 reverse : TAGATCTCTCGAGTCTTGAAGGAC

### Sequences of the host and parasitic RNAs used in the competitive replication assay

**Host-0 (original host RNA)**
GGGAACCCCCCUUCGGGGGGUCACCUCGCGCAGCGGGCUACGCGAGGGAGCCACGCUGCGAAG
CAGCGUGGCGGUUCUCGUGCGUCACCGAAACGCACGAAGGUCGCGCCUCUUCACGAGGCGUC
ACCUGGGAGAGCGCGAAAGCGCUAGCCCGUGCUCUAGCUCUAGAAGGUCUCGAGAUCUCCUC
UAGAGAUAAUUUUGUUUAACUCUAAGAAGGAGAUAUACACAUGCCUAAGACAGCAUCUUCGC
GUAACUCUCUCAGCGCACAAUUGCGCCGAGCCGCGAACACAAGAAUUGAGGCUGAAGGUAAC
CUCGCACUUUCCAUUGCCAACGAUUUACUGUUGGCCUAUGGUCAGUCGCCAUUUAACUCUGA
GGCUGAGUGUAUUUCAUUCAGCCCGAGAUUCGACGGGACCCCGGAUGACUUUAGGAUAAAUU
AUCUUAAAGCCGAGAUCAUGUCGAAGUAUGACGACUUCAGCCUAGGUAUUGAUACCGAAGCU
GUUGCCUGGGAGAAGUUCCUGGCAGCAGAGGCUGAAUGUGCUUUAACGAACGCUCGUCUCUA
UAGGCCUGACUACAGUGAGGAUUUCAAUUUCUCACUGGGCGAGUCAUGUAUACACAUGGCUC
GUAGAAAAAUAGCCAAGCUAAUAGGAGAUGUUCCGUCCGUUGAGGAUAUGUUGCGUCACUGC
CGAUUUUCUGGCGGUGCUACAACAACGAAUAACCGUUCGUACAGUCAUCCGUCCUUCAAGUU
UGCACUUCCGCAAGCGUGUACGCCUCGGGCUUUGAAGUAUGUUUUAGCUCUCAGAGCUUCUA
CACAUUUCGAUAUCAGAAUUUCUGAUAUUAGCCCUUUUAAUAAAGCAGUUACCGUACCUAAG
AACAGUAAGACAGAUCGUUGUAUUGCUAUCGAACCUGGUUGGAAUAUGUUUUUCCAACUGGG
UAUCGGUGGCAUUCUACGCGAUCGGUUGCGUUGCUGGGGUAUCGAUCUGAAUGAUCAGACGA
UAAAUCAGCGCCGCGCUCACGAAGGCUCCGUUACUAAUAACUUAGCAACGGUUGAUCUCUCA
GCGGCAAGCGAUUCUAUGUCUCUUGCCCUCUGUGAGCUCUUAUUGCCCCGAGGCUGGUUUGA
GGUUCUUAUGGACCUCAGAUCACCUAAGGGGCGAUUGCCUGACGGUAGUGUUGUUACCUACG
AGAAGAUUUCUUCUAUGGGUAACGGUUACACAUUCGAGCUCGAGUCGCUUAUUUUUGCUUCU
CUCGCUCGUUCCGUUUGUGAGAUACUGGACUUAGACUCGUCUGAGGUCACUGUUUACGGAGA
CGAUAUUAUUUUACCGUCCCGUGCAGUCCCUGCCCUCCGGGAAGUUUUUAAGUAUGUUGGUU
UUACGACCAAUACUAAAAAGACUUUUUCCGAGGGGCCGUUCAGAGAGUCGUGCGGCAAGCAC
UACUAUUCUGGCGUAGAUGUUACUCCCUUUUACAUACGUCACCGUAUAGUGAGUCCUGCCGA
UUUAAUACUGGUUUUGAAUAACCUAUAUCGGUGGGCCACUAUUGACGGCGUAUGGGAUCCUA
GGGCCCAUUCUGUGUACCUCAAGUAUCGUAAGUUGCUGCCUAAACAGCUGCAACAUAAUACU
AUACCUGACGGUUACGGUGAUGGUGCCCUCGUCGGAUCGGUCCUAAUCAAUCCUUUCGCGAA
AAACCGCGGGUGGAUCCGGUACGUACCGGUGAUUACGGACCAUACAAGGGACCAAGAGCGCG
CUGAGUUGGGGUCGUAUCUCUACGACCUCUUCUCGCGUUGUCUCUCGGAAAGUAACGAUGGG
UUGCCUCUUAGGGGUCCAUCGGGUUGCGAUUUUGCUGAUCUAUUUGCCAUCGAUCAGCUUAU
CUGUAGGAGUGAUCCUACGAAGAUAAGCAGGCCUACCGGUAAAUUCGAUAUACAGUACAUCG

CGUGCUGUUGUUCGGGUUGUUGUUAGGCUUGCGGCCGCACUCGAGAGAUCUAGAGCAUCACG
GUCGAACUCCCGUACGAGGUGCCCGCACCUCGUCCCCCCCUUCCGGGGGGGUCCCC

**Host-99**
GGGAACCCCCCUUCGGGGGGUCACCUCGCGCAGCGGGCUACGCGAGGGGGCCGCGCUGCGAAG
CAGCGUGGCGGUUCACGUGCGUUACCGAAACGCACGAAGGUCGCGCCUCUCCGCGAGGCGUC
ACCUGGGAGAGCGCGAAAGCGCUAGCCCGUGCUCUAGCUCUAGAAGGUCUCGAGAUCUCCUC
UAGAGAUAAUUUUGUUUAACUCUAAGAAGGAGAUAUGCACAUGCCUAAGACAGCAUCUUCGC
GUAACUCUCUCAGCGCACAAUUGCGCCGAGCCGCGAACACAAGAAUUGAGGCUGAAGGUAAC
CUCGCACUUUCCAUUGCCAACGAUUUACUGUUGGCCUAUGGUCAGUCGCCAUUUAACUCUGA
GGCUGAGUGUAUUUCAUUCAGCCCGAGAUUCGACGGGACCCCGGAUGACUUUAGGAUAAAUU
AUCUUAAAGCCGAGAUCAUGUCGAAGUAUGACGACUUCAGCCUAGGUAUUGAUACCGAAGCU
GUUGCCUGGGAGAAGUUCCUGGCAGCAGAGGCUGAAUGUGCUUUAACGAACGCUCGUCUCUA
UAGGCCUGACUACAGUGAGGAUUUCAAUUUCUCACUGUGCGAGUCAUGUAUACACAUGGCUC
GUAGAAAGAUAGCCAAGCUAAUAGGAGAUGUUCCGUCCGUUGAGGAUAUGUUGCGUCACUGC
CGAUUUUCUGGCGGUGCUACAACAACGAAUAACCGUUCGUACAGUCAUCCGUCCUUCAAGUU
UGCGCUUCCGCAAGCGUGUACGCCUCGGGCUUUGAAGUAUGUUUUAGCUCUCAGAGCUUCUA
CACAUUUCGAUAUCAGAAUUUCUGAUAUUAGCCCUUUUAAUGAAGCAGUUACCGUACCUAAG
AACAGUAAGACAGAUCGUUGUAUUGCUAUCGAACCUGGUUGGAAUAUGUUUUUCCAACUGGG
UAUCGGUGGCAUUCUACGCGAUCGGUUGCGUUGCUGGGGUAUCGAUCUGAAUGAUCAGACGA
UAAAUCAGCGCCGCGCUCACGAAGGCUCCGUUACUAAUAACUUAGCAACGGUUGAUCUCUCA
GCGGCAAGCGAUUCUAUGUCUCUUGCCCUCUGUGAGCUCUUAUUGCCCCGAGGCUGGUUUGA
GGUUCUUAUGGACCUCAGAUCACCUAAGGGGCGAUUGCCUGACGGUAGUGUUGUUACCUACG
AGAAGAUUUCUUCUAUGGGUAACGGUUACACAUUCGAGCUCGAGUCGCUUAUUUUUGCUUCU
CUCGCUCGUUCCGUUUGUGAGAUACUGGACUUAGACCCGUCUGAGGUCACUGUUUACGGAGA
CGAUAUUAUUUUACCGUCCCGUGCAGUCCCUGCCCUCCGGGAAGUUUUUAAGUAUGUUGGUU
UUACGACCAAUACUAAAAAGACUUUUUCCGAGGGGCCGUUCAGAGAGUCGUGCGGCAAGCAC
UACUAUUCUGGCGUAGAUGUUACUCCCUUUUACAUACGUCACCGUAUAGUGAGUCCUGCCGA
UUUAAUACUGGUUUUGAAUAACCUAUAUCGGUGGGCCACUAUUGACGGCGUAUGGGAUCCUA
GGGCCCAUUCUGUGUACCGCAAGUAUCGUAAGUUGCUGCCUAAACAGCUGCGACGUAAUACU
AUACCUGACGGUUACGGUGAUGGUGCCCUCGUCGGAUCGGUCCUAAUCAAUCCUUUCGCGAA
AAACCGCGGGUGGAUCCGGUACGUACCGGUGAUUACGGACCAUACAAGGGACCAAGAGCGCG
CUGAGUUGGGGUCGUAUCUCUACGACCUCUUCUCGCGUUGUCUCUCGGAAAGUAACGAUGGG
UUGCCUCUUAGGGGUCCAUCGGGUUGCGAUUUUGCUGAUCUAUUUGCCAUCGAUCAGCUUAU
CUGUAGGAGUGAUCCUACGAAGAUAAGCAGGCCUACCGGUAAAUUCGAUAUACAGUACAUCG
CGUGCUGUUGUUCGGGUUGUUGUUAGGCUUGCGGCCGCACUCGAGAGAUCUAGAGCAUCACG
GUCGAACUCCCGUACGAGGUGCCCGCACCUCGUCCCCCCCUUCCGGGGGGGUCCCC

**Host-115**
GGGAACCCCCCUUCGGGGGGUCACCUCGCGCAGCGGGCUACGCGAGGGAGCCACGCUGCGAAG
CAGCGUGGUGGUUCUCGUGCGUUACCGAAACGCACGAAGGUCGCGCCUCUCCACGAGGCGUC
ACCAUGGGAGAGCGCAAAAGCGCUAGCCCGUGCUCUAGCUCUAGAAGGUCUCGAGAUCUCCU
CUAGAGAUAAUUUUGUUUAACUCUAAGAAGGAGAUAUACACAUGCCUAAGACAGCAUCUUCG
CGUAACUCUUUCAGCGCACAAUUGCGCCGAGCCGCGAACACAAGAAUUGAGGCUGAAGGUAA
CCUCGCACUUUCCAUUGCCAACGAUUUACUGUUGGCCUAUGGUCAGUCGCCAUUUAACUCUG
AGGCUGAGUGUAUUUCAUUCAGCCCGAGAUUCGACGGGACCCCGGAUGACUUUAGGAUAAAU
UAUCUUAAAGCCGAGAUCAUGUCGAAGUAUGACGACUUCAGCCUAGGUAUUGAUACCGAAGC

UGCUGCCUGGGAGAAGUUCCUGGCAGCAGAGGCUGAAUGUGCUUUAACGAACGCUCGUCUCU
AUAGACCUGACUACAGUGAGGAUUUCAAUUUCUCACUGGGCGAGUCAUGUAUACACAUGGCU
CGUCGAAAAAUAGCCAAGCUAAUAGGAGAUGUUCCGUCCGUUGAGGAUAUGUUGCGUCACUG
CCGAUUUUCUGGCGGUGCUACAACAACGAAUAACCGUUCGUACAGUCAUCCGUCCUUCAAGU
UUGCACUUCCGCAAGCGUGUACGCCUCGGGCUUUGAAGUAUGUUUUAGCUCUCAGAGCUUCU
ACACAUUUCGAUAUCAGAAUUUCUGAUAUUAGCCCUUUUAAUGAAGCAGUUACCGUACCUAA
GAACAGUAAGACAGAUCGUUGUAUUGCUAUCGAACCUGGUUGGAAUAUGUUUUUCCAACUGG
GUAUCGGUGGCAUUCUACGCGAUCGGUUGCGUUGCUGGGGUAUCGAUCUGAAUGAUCAGACG
AUAAAUCAGCGCCGCGCUCACGAAGGCUCCGUUACUAAUAACUUAGCAACGGUUGAUCUCUC
AGCGGCAAGCGAUUCUAUGUCUCUUGCCCUCUGUGAGCUCUUAUUGCCCCGAGGCUGGUUUG
AGGUUCUUAUGGACCUCAGAUCACCUAAGGGGCGAUUGCCUGACGGUAGUGUUGUUACCUAC
GAGAAGAUUUCUUCUAUGGGUAACGGUUACACAUUCGAGCUCGAGUCGCUUAUUUUUGCUUC
UCUCGCUCGUUCCGUUUGUGAGAUACUGGACUUAGACUCGUCUGAGGUCACUGUUUACGGAG
ACGAUAUUAUUUUACCGUCCCGUGCAGUCCCUGCCCUCCGGGAAGUUUUUAAGUAUGUUGGU
UUUACGACCAAUACUAAAAAGACUUUUUCCGAGGGGCCGUUCAGAGAGUCGUGCGGCAAGCA
CUACUAUUCUGGCGUAGAUGUUACUCCCUUUUACAUACGUCACCGUAUAGUGAGUCCUGCCG
AUUUAAUACUGGUUUUGAAUAACCUAUAUCGGUGGGCCACUAUUGACGGCGUAUGGGAUCCU
AGGGCCCAUUCUGUGCACCUCAAGUAUCGUAAGUUGCUGCCUAAACAGCUGCAACAUAAUAC
UAUACCUGACGGUUACGGUGAUGGUGCCCUCGUCGGAUCGGUCCUAAUCAAUCCUUUCGCGA
AAAACCGCGGGUGGAUCCGGUACGUACCGGUGAUUACGGACCAUACAAGGGACCAAGAGCGC
GCUGAGUUGGGGUCGUAUCUCUACGACCUCUUCUCGCGUUGUCUCUCGGAAAGUAACGAUGG
GUUGCCUCUUAGGGGUCCAUCGGGUUGCGAUUUUGCUGAUCUAUUUGCCAUCGAUCAGCUUA
UCUGUAGGAGUGAUCCUACGAAGAUAAGCAGGCCUACCGGUAAAUUCGAUAUACAGUACAUC
GCGUGCUGUUGUUCGGGUUGUUGUUAGGCUUGCGGCCGCACUCGAGAGAUCUAGAGCAUCAC
GGUCGAACUCCCGUACGAGGUGCCCGCACCUCGUCCCCCCCUUCCGGGGGGGUCCCC

**Parasite-α13**
GGGAACCCCCCUUCGGGGGGUCACCUCGCGCAGCGGGCUGCGCGAAGGAGCCACGCUGCGAAG
CAGUGUGGCGGUUCUCGUGCGUUACCGAAACGCACGAAGGUCGCGCCUCUUCACGAGGCGUC
ACCUGGGAGAGCGCGAAAGCGCUAGCCCGUGAUUCGUCACGGUCGAACUCCCGUACGAGGUG
CCCGCACCUCGUCCCCCCCUUCCGGGGGGGUCCCC

**Parasite-β99**
GGGAACCCCCCUUCGGGGGGUCACCUCGCGCAGCGGGCUACGCGAGGGAGCCACGCUGCGAAG
CAGCGUGGCGGUUCUCGUGCGUUACCGAAACGCACGAAGGUCGCGCCUCUCCGCGAGGCGUC
ACCUGGGAGAGCGCGAAAGCGCUAGCCCGUGCUCUAGCUCUAGAAGGUCUCGAGAUCUCCUU
UAGAGAUAAUUUUGUUUAACUCUAAGAAGGAGAUAUGCACAUGCCUAAGACAGCAUCUUCGC
GUAACUCUUUCAGCGCACAAUUGCGCCGAGCCGCGAACACAAGAAUUGAGGCUGAAGGUAAC
CUCGCACUUUCCAUUGCCAACGAUUUACUGUUGGCCUAUGGUCAGUCGCCAUUUAACUCUGA
GGCUGAGUGUAUUUCAUUCAGCCCGAGAUUCGACGGGACCCCGGAUGACUUUAGGAUAAAUU
GUCUUAAAGCCGAGGUCAUGUCGAAGUAUGACGACUUCAGCCUAGGUAUUGAUACCGAAGCU
GUUGCCUGGGAGAAGUUCCUGGCAGCAGAGGCUGAAUGUGCUUUAACGAACGCUCGUCUCUA
UAGGCCUGACUACAGUGAGGAUUUCAAUUUCUCACUGGGCGAGUCAUGUAUACACAUGGCUC
GUGGAAAAAUAGCCAAGCUAAUAGGAGAUGUUCCGUCCGUUGAGGAUAUGUUGCGUCACUGC
CGAUUUUCUGGCGGUGCUACAACAACGAAUAACCGUUCGUACAGUCAUCCGUCCUUCAAGUU
UGCACUUCCGCAAACGUGUACGCCUCGGGCUUUGAAGUAUGUUUUAGCUCUCAGAGCUUCUA
CACAUUUCGAUAUCAGAAUUUCUGAUAUUAGCCCUUUUAAUGGGUUGCGAUUUUGCUGAUCU

AUUUGCCAUCGAUCAGCUUAUCUGUAGGAGUGAUCCUACGAAGAUAAGCAGGCCUACCGGUA
AAUUCGAUAUACAGUACAUCGCGUGCUGUUGUUCGGGUUGUUGUUAGGCUUGCGGCCGCACU
CGAGAGAUCUAGAGCAUCACGGUCGAACUCCCGUACGAGGUGCCCGCACCUCGUCCCCCCCUU
CCGGGGGGGUCCCC

**Parasite-γ115**
GGGAACCCCCCUUCGGGGGGUCACCUCGCGCAGCGGGCUACGCGAGGGAGCCACCUGCGAAGC
AGCGUGGCGGUUCUCGUGCGUUACCGAAACGCACGAAGGUCGCGCCUCUCCACGAGGCGUCA
CCAUGGGAGAGCGCAAAAGCGCUAGCCCGUGCUCUAGCUCUAGAAGGUCUCGAGAUCUCCUC
UAGAGAUAAUUUUGUUUAACUCUAAGAAGGAGAUAUGCUUUAACGAACGCUCGUCUCUAUAG
ACCUGACUACAGUGAGGAUUUCAAUUUCUCACUGGGCGAGUCAUGUAUACACAUGGCUCGUC
GAAAAAUAGCCAAGCUAAUAGGAGAUGUUCCGUCCGUUGAGGAUAUGUUGCGUCACUGCCGA
UUUCCUGGCGGUGCUACAACAACGAAUAACCGUUCGUACAGUCAUCCGUCCUUCAAGACUCG
AGAGAUCUAGAGCAUCACGGUCGAACUCCCGUACGAGGUGCCCGCACCUCGUCCCCCCCUUCC
GGGGGGGUCCCC
